# Eyes on nature: Embedded vision cameras for multidisciplinary biodiversity monitoring

**DOI:** 10.1101/2023.07.26.550656

**Authors:** Kevin F.A. Darras, Marcel Balle, Wenxiu Xu, Yang Yan, Vincent G. Zakka, Manuel Toledo-Hernández, Dong Sheng, Wei Lin, Boyu Zhang, Zhenzhong Lan, Li Fupeng, Thomas C. Wanger

## Abstract

Global environmental challenges require comprehensive data to manage and protect biodiversity. Currently, vision-based biodiversity monitoring efforts are mixed, incomplete, human-dependent, and passive. To tackle these issues, we present a portable, modular, low-power device with embedded vision for biodiversity monitoring. Our camera uses interchangeable lenses to resolve barely visible and remote subjects, as well as customisable algorithms for blob detection, region-of-interest classification, and object detection to identify targets. We showcase our system in six case studies from the ethology, landscape ecology, agronomy, pollination ecology, conservation biology, and phenology disciplines. Using the same devices, we discovered bats feeding on durian tree flowers, monitored flying bats and their insect prey, identified nocturnal insect pests in paddy fields, detected bees visiting rapeseed crop flowers, triggered real-time alerts for waterbirds, and tracked flower phenology over months. We measured classification accuracies between 55% and 96% in our field surveys and used them to standardise observations over highly-resolved time scales. The cameras are amenable to situations where automated vision-based monitoring is required off the grid, in natural and agricultural ecosystems, and in particular for quantifying species interactions. Embedded vision devices such as this will help addressing global biodiversity challenges and facilitate a technology-aided global food systems transformation.

## Main text

### Introduction

This century’s rapid biosphere transformations exert formidable pressure on nature and people^1,2^: land use conversion is undiminished, climate change is accelerating, infectious disease risks are increasing, food systems are reaching maximal capacity, and disturbance of remaining natural ecosystems is higher than ever. These pressures are closely linked to biodiversity, both an actor providing vital services to humans and their ecosystems, and a victim of these developments3. To prevent, manage and mitigate these global transformations, we need high resolution biodiversity data^4,5^. We are heavily reliant on artificial intelligence to analyse the resulting big data and to tackle Sustainable Development Goals ^6–8^. However, while climate and land use change are internationally monitored with remote sensing and coordinated research networks, and unlike biotechnologies that are powered by modern laboratory methods and artificial intelligence^9–12^, much progress has yet to be made at intermediate spatial scales towards the automated monitoring of biodiversity^13^.

Much of biodiversity is sampled with vision, and we are facing critical challenges to attain comprehensive monitoring. First, heterogenous methods and scattered data sources for different taxa, biomes, and disciplines are hampering progress on integrated analyses^14–17^. Cultural challenges slow their progress even though frameworks and infrastructures have been proposed^18,19^. Second, sampling coverage and resolution along spatial and temporal scales are insufficient, resulting in considerable knowledge biases and gaps^15,20–22^ that prevent effective conservation: notably, most of the challenging animals to monitor (amphibians, insects, mammals, and reptiles) are data-deficient and likely threatened^23^. Third, many methods still depend on human labour, and are thus time consuming, error-prone, and hard to reproduce^24–26^, a problem further compounded downstream by the scarcity of taxonomic experts^27^. Fourth and lastly, most passive vision-based methods lack real-time feedback^28^, thus ruling out immediate interventions despite increased uptake of adaptive management approaches^29,30^. Theoretically, these challenges may be solved by deploying continuously-powered digital imaging devices for sampling multiple taxa across large scales, with embedded artificial intelligence to process data in real-time at the edge and to trigger meaningful reactions when needed. While the underlying technologies exist and open hardware abounds^31^, devices are still in early development stages and we lack integrated solutions^32,33^.

We harnessed recent technological advances into the first field-ready, portable, low-power, modular, embedded vision camera system - dubbed “ecoEye” - thus taking advantage of recent progress in embedded computing^34^ and also achieving set goals for high-resolution, long-term, real-time, and standardised sampling methods^13^. Our system can: 1) non-invasively monitor various taxa for different applications across disciplines; 2) reach high temporal resolution and coverage with solar power and be scaled up in space due to its moderate cost and size; 3) analyse images in real-time using established computer vision algorithms and performance assessment workflows that standardise observation results, and thus 4) link specific detections to real-time reactions for intervening in environmental processes. We finally discuss how our and other embedded vision devices fare for addressing key vision-based biodiversity monitoring challenges.

### Results

We conducted six case studies representing different use cases and disciplines in several regions of China (Tab. 1, Fig. 1). We deployed our cameras with setups and scripts adjusted for various day- and nighttimes and different target organisms. Both frame differencing-based blob detection as well as deep learning convolutional neural networks (CNNs hereafter) based on MobileNet-v2^35^ and trained with images captured by the cameras during test deployments were used to detect and identify targets in real-time on the cameras during the actual survey deployments. The performance of the detection and classification algorithms were evaluated with survey images; we report them as field accuracies (i.e., F1-scores).

**Figure 1:**
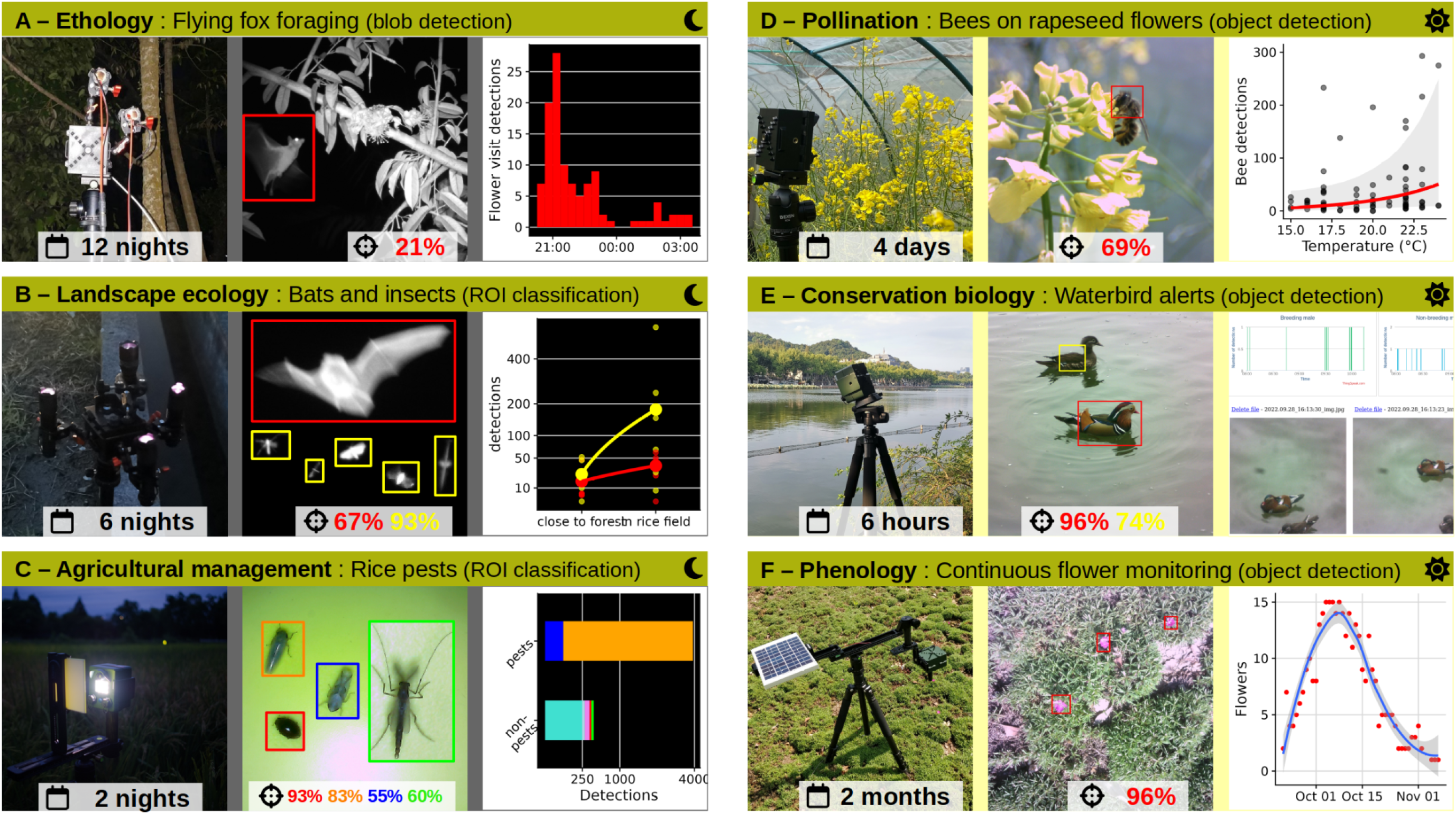
Field-tested applications for the embedded vision camera. Middle panels for B) and C) were made from multiple pictures to show the diverse target types. Padding was added to the bounding boxes of the detections for better visibility of the targets. The crosshairs represent field accuracies, measured with F1 scores.

**Figure 2:**
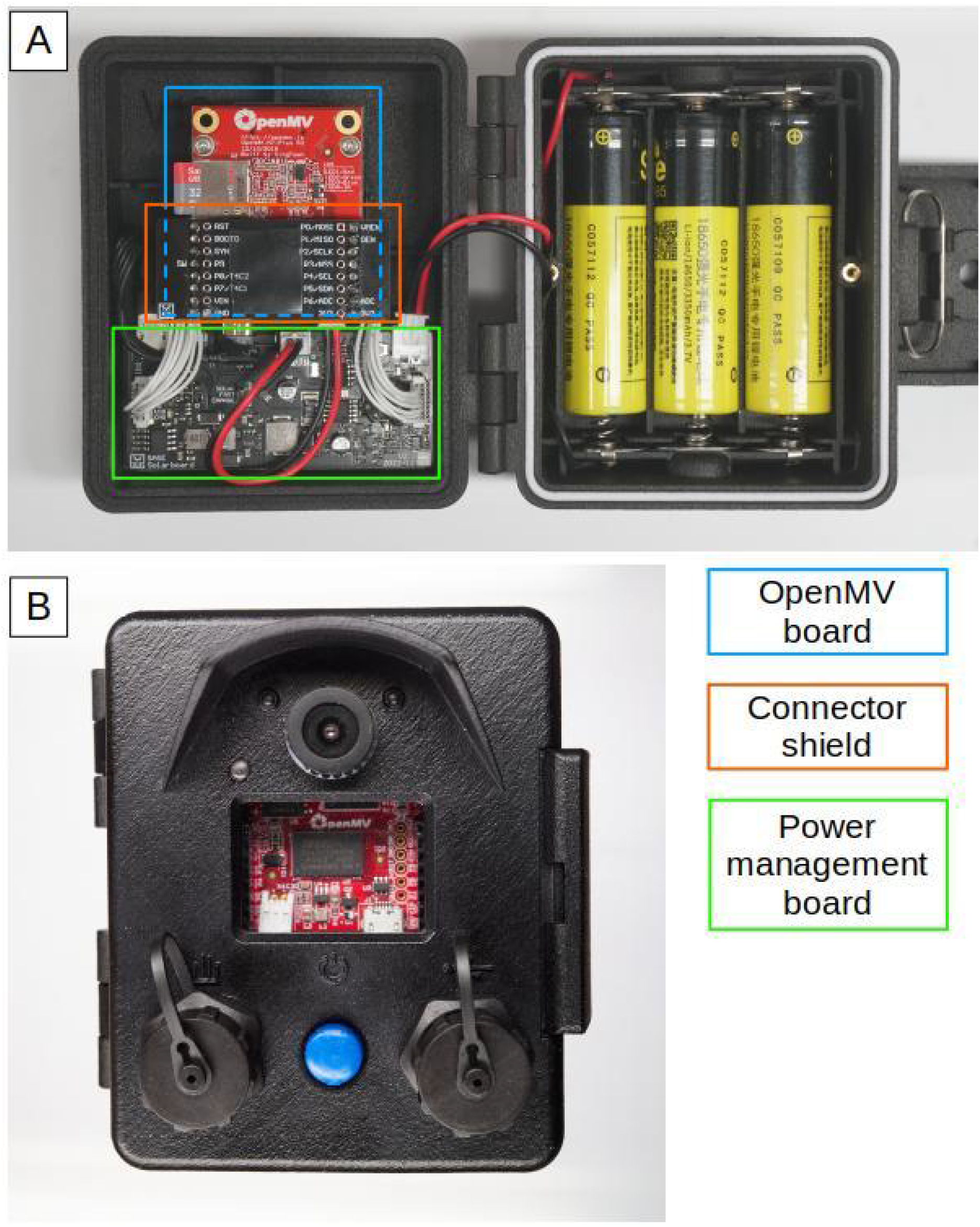
Inner (A) and outer front (B) views of the “ecoEye” camera

**Table 1:**
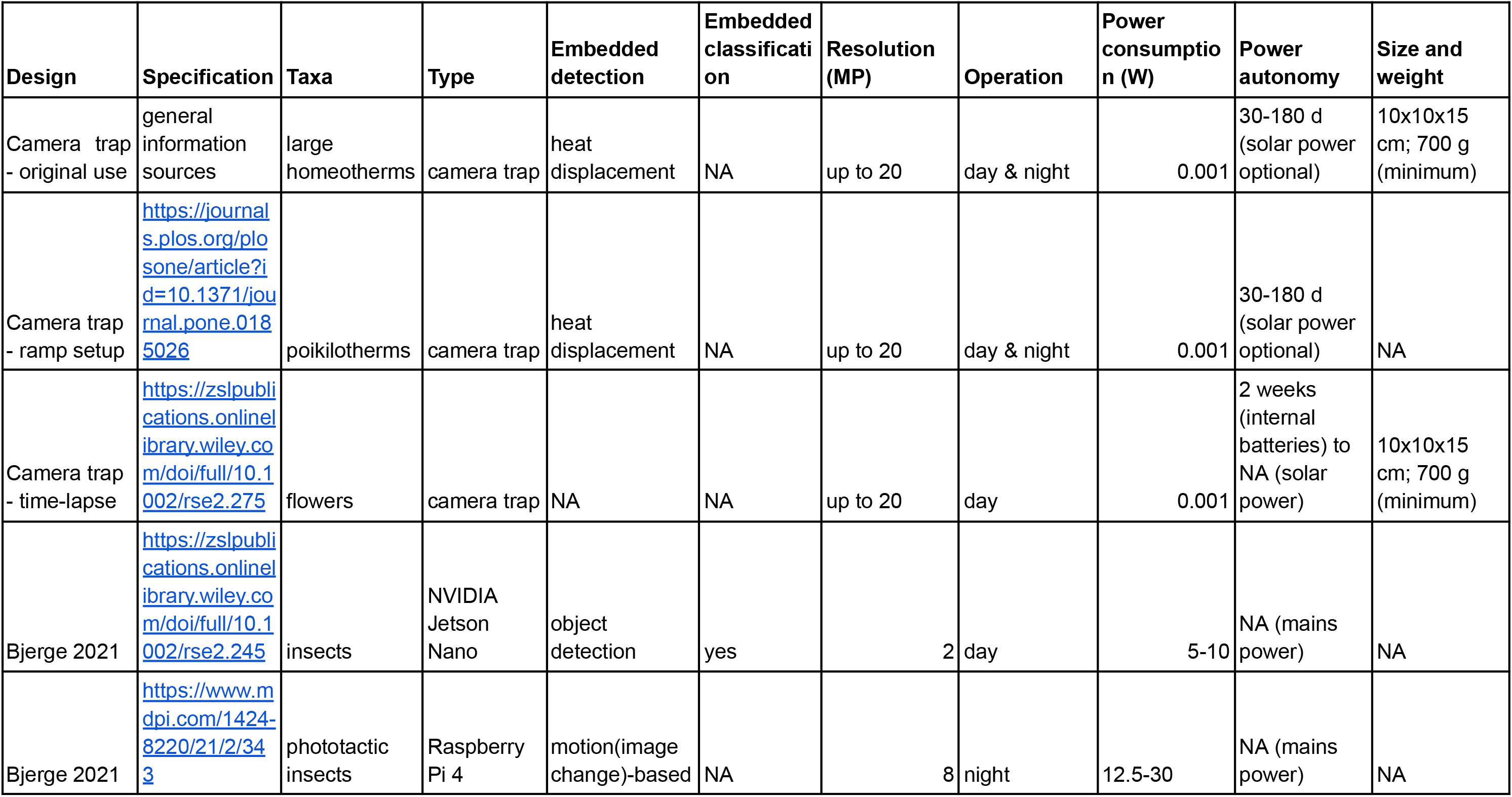
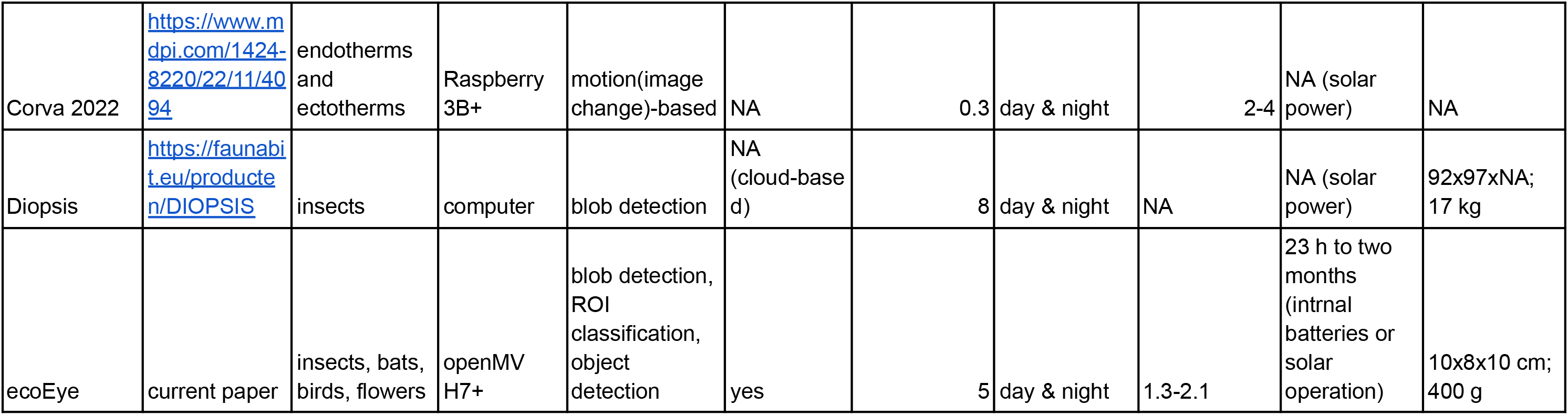
Comparison of automated vision-based field biodiversity monitoring devices. The size and weight indications exclude support and continuous powering accessories.

#### A) Behavioural Ecology: Monitoring bats visiting Durian tree flowers

Bats provide important pollination services for tropical crop trees but little is known about their foraging behavior^36^ and current vision-based monitoring approaches are a compromise between automation and resolution^37,38^. In Southeast Asia, the Durian tree produces the “king of fruits”, whose most common pollinators are fruit bats^39^. On the island of Hainan, in South China, plantations were established around 1950^40^, and flowers were usually pollinated by hand, although natural pollination seems to be occurring too^39^ (pers. comm. LF). Although fruit bats inhabit the island, they have never been reported to pollinate Durian there. We therefore surveilled Durian flower visitors in Hainan, China (Fig. 1A) by monitoring the nocturnal behaviour of unknown targets at high resolution, using near-infrared illumination and blob detection algorithms, from the ground.

We monitored one flowering tree for 12 nights at an average of 0.68 (min: 0.56; max: 0.89) frames per second with blob detection to detect image changes above a threshold area and color deviation (Fig 1A). We manually screened 23 154 triggered images and found the flying fox *Roussettus leschenaultii* in 122 images. The manual screening allowed us to determine optimal detection parameters (minimum blob area range: 95 000 - 135 000 and maximum blob area range: 185 000 - 475 000) for each deployment date, thus obtaining a mean maximum accuracy of 0.21 over deployment nights (range: 0.10-0.34). Bats were detected flying towards or from flowers in 63, and feeding on flowers in 59 images. Flowers were visited 9 times per night on average (range: 3-32 visits, N=7; flowers open only for one night). Two activity peaks were evident: early in the night (∼21:00) and shortly before dawn twilight (∼3:00).

#### B) Landscape ecology: Determining bat and insect occurrence in rice fields

Landscape composition determines the occurrence of mobile predators and their prey. Insectivorous bats, which are sensitive to forest edges^41^ and waterways^42^, regulate insect populations in natural and agricultural landscapes through predation, thus providing, and biocontrol services. Rice in particular benefits from economically valuable pest control services^43,44^. Current passive sampling methods use digital imagery to sample only insects or bats^38,45^, so we used our cameras with region-of-interest classification and near-infrared illumination to remotely detect and identify both these differently-sized nocturnal targets.

We monitored flying insects and bats in a mixed rice paddy and forest landscape for 6 nights in summer of 2022 during astronomical dusk (between approximately 19:20 and 20:30), in Hangzhou, China (Fig. 1B). Each night, we set up one camera inside the rice field and one at its border, adjacent to a waterway with a forest on its other shore, and pointed them to the night sky. The cameras were operating with a sensitive blob detection threshold at an average of 1.32 (range: 0.37-1.88) frames per second. Flying objects were detected as blobs up to an estimated 7 m above ground, and images extracted from their bounding boxes were classified into bats, insects, or unknown flying objects with our trained CNN. We obtained a maximum field classification accuracy (F-score) of 67% for bats and 93% for insects at their individual optimal confidence thresholds of 0.6 and 0.3. After considering only detections above these thresholds, we obtained a nightly average of 16 bat and 25 insect detections close to the forest compared to 37 bat and 180 insect detections inside the rice field, with a significantly lower bat to insect detections ratio inside the rice field than close to the forest (Estimate = -0.34 ±0.17 SD, Probability_(Estimate=0 | Null hypothesis)_ = 0.04).

#### C) Agricultural sciences: Quantifying nocturnal insect rice pests

Monitoring pests is essential for agricultural management. Light traps are a common sampling method for flying insect pests in rice paddy fields^46^, a staple food throughout Asian countries that is damaged by the brown and white-backed planthoppers. Existing monitoring devices are bulky, costly, sometimes lethal and have no embedded classification^47,48^. We used our cameras to demonstrate an embedded multiple classification task of nocturnal insects attracted to non-lethal, portable light traps, with real-time cloud transmission of results.

We monitored nocturnal, flying, phototaxic insects attracted to white-light illuminated plastic boards. Cameras were installed in two locations inside a rice field in Hangzhou, China (Fig. 1C) and worked for two hours on two nights in 2022 and 2023, starting at nautical dusk, around 17:30. Cameras operated with a sensitive blob detection threshold at an average of 0.41 (min: 0.40; max: 0.41) frames per second. Detection numbers for each detected target were sent to the cloud, via an internal plug-in WLAN module connected to a 4G portable router, with a transfer success rate between 76% and 83%. Landing objects were blob-detected and their bounding boxes extracted for classification in nine classes with our trained CNN. For the different morphospecies detected during survey deployments, we obtained maximum field accuracies ranging from 8% (Curculionidae - only 39 training images) to 93% (Coleoptera) for non-pest species (mean: 53%, median: 60%), and from 55% to 83% for potential planthopper pests (two Delphacidae morphospecies). After retaining only detections above the corresponding optimal confidence thresholds, we found 3856 detections for the Delphacidae 1 and 59 detections for the Delphacidae 2 morphospecies, identified as the brown (*Nilaparvata lugens*) and white-backed (*Sogatella furcifera*) planthoppers respectively, and 452 non-pest detections across both survey nights.

#### D) Pollination ecology: Monitoring bees on rapeseed flowers

Global insect pollinator declines are disrupting pollination networks and crop production^49^, and monitoring is essential to reap their benefits^50^. Long-term diel monitoring of bees was attempted with motion-detection devices, albeit with complicated setups^51^. We used our cameras to monitor rapeseed, a major oil crop, with object detection models specifically trained for identifying pollination events by bees.

We monitored solitary bees (*Osmia bicornis*) on rapeseed flowers growing inside experimental enclosures with six camera deployments over four summer days in Fuyang district, China (Fig. 1D). Targets were searched in each image with our object detection CNN. We operated the cameras with a confidence threshold of 0.5 at an average of 1.37 (range: 1.13-1.91) frames per second, and obtained an average field classification accuracy of 69% (range: 55-89%) over all deployments (Fig. S4). As field accuracy varied among deployments, we standardised the data to obtain true detection counts, which we used to derive temporal bee activity profiles that were comparable across deployments (Fig. S4). We evidenced a highly probable positive relationship between standardized bee detections and ambient temperature (Estimate = 0.24, Probability_(Estimate>0)_ = 0.97), in line with previous studies^51^.

#### E) Conservation Biology: Protected waterbird real-time alerts

Adaptive management of habitats and species is an attractive solution for dealing with unpredictable global changes that precipitate biodiversity loss^29^. Real-time monitoring of protected or endangered species, which typically occur sporadically and in small numbers, is required over long time ranges without human disturbance. Quick reactions to population changes^52^ are possible to potentially initiate fast-response management actions. Birds, as fast-moving, small targets are especially challenging to monitor automatically with the current technology^53,54^. Here, we show how cameras can be networked to send real-time cloud alerts upon detection of specific waterbird species in a protected area.

We monitored swimming mandarin ducks (*Aix galericulata*) using a camera in a nature reserve, from the shore of the Xihu lake in Hangzhou, China, over 6 hours (Fig. 1E). The camera executed an object detection model on each image with a conservative confidence threshold of 0.5, and ran at 1.76 frames per second. It transferred images containing mandarin duck detections exceeding a confidence threshold of 0.5 via an internal plug-in WLAN module connected to a 4G portable router, to a cloud server where the detection time graphs and downsampled images could be checked in real-time (Fig. 1E). 82% of the detections data and 64% of the images were successfully transferred from the cameras to the server. We obtained maximum field accuracies of 74% for females, and 96% for breeding males at their optimal confidence thresholds, which allowed to filter the dataset to a total of 594 raw detections, yielding 134 female and 405 male standardised detections after correcting for individual detection accuracies.

#### F) Phenology: Flowering plants

Phenology cameras are used to monitor vegetation phenology when high temporal resolution and long sampling durations are needed for instance to understand climate change impacts. However, these cameras are not able to analyse image contents, and even though automated analysis could happen post-capture^55^, new monitoring methods are required for greater reactivity^56^. We show here how our cameras can operate over long periods for monitoring the phenology of inanimate, non-animal targets such as flowering plants with high accuracy.

We monitored ground-dwelling Carthusian pink (*Dianthus carthusianorum*) flowers with two cameras situated on our university campus in Hangzhou, China, over a period of two months (Fig. 1F). The cameras continuously captured images at an interval of 15 minutes from the start until the end of civil twilight (approximately 5:30 to 18:00), as they were automatically recharged with 5 W solar panels during sunny weather. The cameras ran an object detection model at a conservative confidence threshold of 0.1 to detect multiple flowers within each image, and all images were saved. We attained a maximum field accuracy of 96% for detecting open Carthusian pink flowers. We automatically tracked single flowers over frames and days using time series clustering in R^57^ to establish a flowering peak on October 5, 2022 with 15 simultaneously open flowers and a mean lifespan of single flowers of 4 days (range: 0.4-13.5 d).

### Discussion

Embedded vision devices such as the one proposed here have far-reaching implications for environmental research. We demonstrated the potential of our ecoEye cameras for biodiversity monitoring such as phenological observation, real-time species surveillance, and for key ecological interactions such as pollination, pest prevalence and potential pest control. Our versatile design can be utilised for multiple taxa and scaled-up to large spatial and temporal extents due to its interchangeable lens options, sufficient frame rates and resolution, and low power consumption compared to alternative devices (Tab. 1).

The interchangeable lens design with adjustable focus enabled the largest taxonomic coverage and deployment flexibility compared to alternative devices: from millimeter-scale insects, to remote flying bats and flowers on the ground (Table 1). Alternative imaging devices without embedded vision can be used to detect insects^45,58–60^, and recent designs with embedded vision integrate motion-detection^47^ or species classification^61^, but stay confined to their integrated lenses’ angle of view and the corresponding targets that can be imaged. An interchangeable lens and sensor design also allows near-infrared imaging, as cut-off filters can be removed to detect nocturnal organisms lit with near-infrared light, which is invisible to most animals^62^. We attached up to 4 torchlights for lighting remote or fast objects, enabling non-intrusive monitoring of bats or flowers without interference from phototactic insects. While camera traps and the PICT camera also offer near-infrared imaging as an option, none of the embedded vision alternatives currently do (Table 1). Furthermore, the imaging sensors can be swapped for thermal imaging modules for a potentially very effective detection of homeotherms^63^. In comparison, established camera traps have a limited angle of view by design, and can only detect heat displacement of medium to large homeotherms. They have been adapted with poor performance or excessive efforts beyond their designed application: for far-away birds, risky arboreal sampling, nocturnal flying insect traces, with poikilothermic animal ramps, or for inanimate plants^55,64–68^. This underlines the necessity for broader image-based, embedded analytical approaches such as ours, which could even be extended to microscopic scales to monitor zooplankton^69^.

The ecoEye camera is able to sample a large temporal and spatial coverage. Our cameras operated between 0.4 and 1.7 frames per second depending on the set resolution and used algorithm (i.e., between 2.3 and 12.6 MB per second). While frame rates are higher at lower resolutions, this already sufficed for monitoring short-lived pollination events or bat passes. In contrast, camera traps may miss quickly-passing and stationary targets by design^70^. However, all currently available cameras compared here offer high sampling resolutions over coverage times that exceed human capabilities (Table 1), and while other embedded vision devices running on mains power could theoretically be solar-powered, only ours demonstrated a successful implementation over months. Alternative embedded vision devices are also currently bulkier and more expensive (Table 1), so that coverage may not be easily scaled-up in space. Similarly cheap and compact devices such as the PICT cameras or camera traps can also reach high spatial coverage^71^ through replication, but come with challenges in post-deployment data processing demands due to the lack of embedded vision. In the end, the low cost and small size of our devices facilitates covering large spatial scales.

We used standard Artificial Intelligence (AI) evaluation procedures to measure algorithm accuracies in the field and obtain standardised biodiversity data. Vision-based biodiversity detection processes are rarely standardised^72^. AI drives global camera trap syntheses^73^, but the data are derived from a separate sampling process that remains problematically unquantifiable or hard to estimate with ground-truth^74,75^. We measured the accuracies of our algorithms by screening every saved frame over a calibration period for true and false positives and negatives to infer “field accuracies” and mathematically derive standardised detection numbers. Similar calibration approaches were employed with embedded vision devices^61^. In our case, the same CNN’s performance could be corrected across deployments of the pollination ecology use case. We argue that since the realised performance of custom-trained CNNs is inevitably variable, standardising results with field data is necessary and cost-effective at scale - probably also for fine-tuned models based on global datasets^76,77^. As a result, we can infer the true number of events, circumvent complex analytical modeling approaches, and potentially derive densities over the cameras’ field-of-views. We could thereby potentially harmonise sampling methods across realms and taxa. Such standardised data should facilitate large-scale syntheses based on field studies with variable setups at scale, yielding actionable density-based evidence for biodiversity management plans.

Embedded vision is not limited to passive biodiversity monitoring and extends to real-time triggered reactions. At its simplest, the embedded design accelerates the scientific workflow by providing readily-usable detections data with defined metrics, such as the blob characteristics or classification probabilities for pests or pollinators in our use cases. Naive data from blob detection workflows - that are needed in every situation where training images are not available - may be sent to cloud servers for post-classificatio with advanced deep learning models that are less power- and computoation-constrained. Beyond this, data above a threshold confidence level can be used to trigger critical alerts, as we sent and aggregated data in real-time to trigger alerts on cloud servers for pest management or species protection. In comparison, only one of the alternative embedded vision devices was networked for real-time data transmission^61^. Although devices without embedded vision such as “cellular” camera traps can also be networked, costs for sending prevalently irrelevant images can be prohibitively high, justifying “edge” solutions^78^, even though hybrid solutions exist^28,79^. Real-time reactions matter most for remote locations and time-sensitive applications: bat and insect detections can trigger alerts when biocontrol rates fall below levels that are safe for crops; suction samplers could catch flower visitors for DNA barcoding; insecticide spraying or electrocution could be activated upon detection of specific pests on light boards; finally, poacher detections could direct law enforcement or activate deterrents. Furthermore, the core of our system can be mounted on drones to extend spatial coverage and dynamically approach targets^80–82^. Our embedded vision cameras thus offer unprecedented opportunities for triggering meaningful actions.

Embedded vision devices are poised to dethrone camera traps, currently the gold standard for vision-based monitoring, whose various constraints prevent broad scale implementation^83^. Future technological innovations such as ever-improving object detection models, on-sensor machine learning, ultra-low power chips, higher battery energy densities, and tiny machine learning will tip the scale towards higher-resolution imaging and more sophisticated analysis^84^. Open-set recognition will help to tackle the detection of taxonomically unresolved taxa^85^, and individual animals will become distinguishable^86^. Beyond this, the monitoring data should be used for predicting future trends^13^, and technology-enabled adaptive management of ecosystems can be envisaged. However, just like new observation technologies have historically driven scientific progress, the sheer possibilities of embedded vision systems necessitate regulation to alleviate ethics, privacy, and security issues. Likewise, harmonisation efforts should be pursued on a much higher level by devising official standards for computer-vision based monitoring of biodiversity, as to enable truly interoperable data for large syntheses. We hope that with tools that become ever more efficient, we will be able to address challenges faced by nature and society and focus on the implementation of solutions.

## Methods

### Camera design and setup

Our embedded vision camera consists of 1) a low-power, expandable microcontroller board with a modular image sensor; 2) a data transfer and power management board; and 3) a waterproof housing for the boards, batteries, lens mount, and accessories.

The microcontroller board (openMV H7+ computer vision board) is based on the STMicroelectronics STM32H7 microcontroller and handles imaging and data processing from the plug-in imaging board, based on the 1/4” (3.6 x 2.7 mm) Omnivision OV5645 (5 megapixel RGB CMOS sensor). The board is programmed with the open-source openMV Integrated Development Environment (IDE) software in micropython language. The board can be used with other plug-in imaging boards (far-infrared, global shutter RGB), and has separate connectors for connectivity (WLAN) or lighting (white LED) modules, among others.

We developed a power management board based on USB (Injoinic Technology, IP2312) and solar (Consonance Electronics, CN3791) chargers, as well as step-up converter (Aerosemi, MT3608L) integrated circuits; these components control powering the microcontroller board and recharging the internal batteries with external power sources such as from USB or solar panels. Waterproof ports transfer data from USB or external sensors to the microcontroller board. A waterproof push button on the exterior, combined with a soft latch switch circuit, enables manual power-on and safe shutdown of the main board through software. Lastly, a PCF8563 real-time clock chip (NXP Semiconductors) is used for time keeping and pre-scheduled system power-on by the on-chip timer interrupt output.

The housing secures the internal components against mechanical shocks, water ingress, and theft with a rubber gasket, latch, and optionally steel cables and locking screws. It is resistant to immersion (up to 60 seconds, 10 cm depth from front face) and enables waterproof mounting of M12 interchangeable lenses. Up to three 18650 lithium (Li-ion or Li-Po) batteries can be inserted in parallel connection. There is space for two expansion modules plugged into 2.54 mm pitch, 8-pin headers at the front (e.g., LED module) and back (e.g., WLAN module) of the main board. Five mounting points (on all faces except the front) with standard UNC ¼-20 threaded inserts allow mounting accessoires such as lights and tripods. The housing has a transparent window for light from internal plug-in LED boards.

Prior to field deployments, we attached lenses with appropriate angle of view and aperture to the camera, and inserted three 3350 mAh batteries per camera for a total capacity of 10050 mAh. A field deployment consists of six setup steps. First, the camera is installed at the study site, pointing at the target. Second, the camera is connected with a laptop; we wrote custom micropython scripts to control the microcontroller board through the openMV IDE. Third, we run the script in live view mode to adjust the image composition and focus the lens by screwing it into the threaded lens mount. Fourth, we run the script in test mode to check the output of our chosen detection settings in the terminal. Fifth, the script with the chosen parameters is saved on the camera, which is disconnected from the computer and then powered on with the external power button to execute the script. Lastly, proper camera operation can be checked with LED signals corresponding to pre-programmed events. The cameras can be set up for saving all pictures, only triggered pictures, or none, while saving image and detection logs as CSV files. An overview of setup parameters is given in Table 2.

**Table 2:**
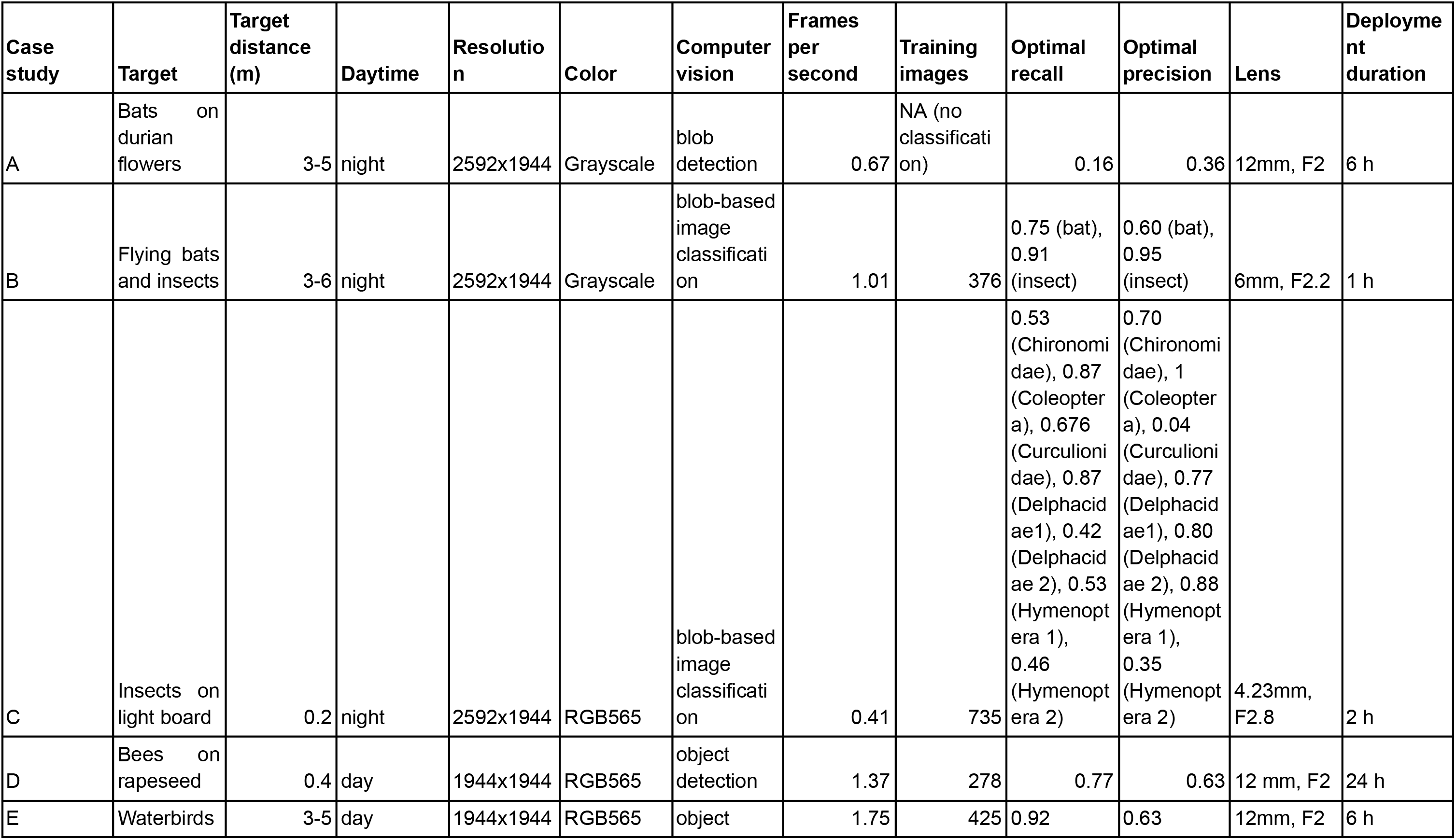
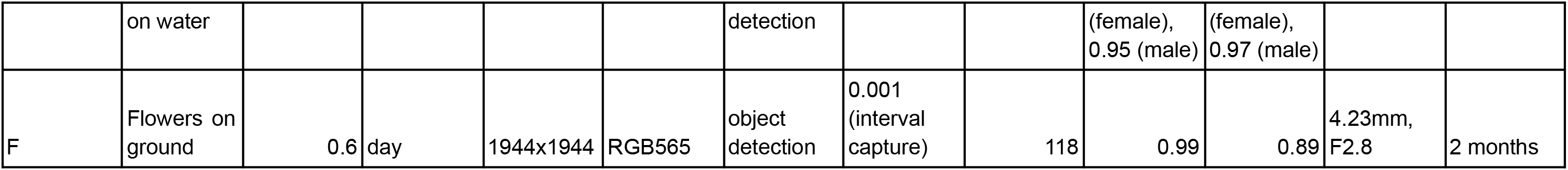
Comparison of case study setups.

### Embedded vision

When images for training CNNs were not available, we used frame differencing to detect changes in the image as “blobs”^87^. Changes are obtained from computing the difference between the current and a reference image. We adapted to changing image conditions by regularly updating the reference image (every minute) by blending the current image into the reference image by half (alpha compositing value of 128 out of 256), but only when no image change was detected (maximum extension by one minute). When we applied automatic exposure time and image gain to adjust to variable lighting conditions, the exposure and gain adjustments were performed shortly before and fixed until the next reference image update. We restricted the range of pertinent blobs by choosing lower and sometimes upper blob size limits (in pixels) and determined the detection algorithm’s colour sensitivity by choosing minimum colour thresholds that need to be exceeded (in Lab or grayscale colour spaces) by the blobs. We used blob detection primarily with controlled image backgrounds to avoid false positives under changing conditions.

When enough training images were available, we explicitly located multiple targets with CNNs for region of interest classification or for object detection. These algorithms enable the detection of targets that have too small sizes for whole-image classification methods. For region-of-interest classification, we classified image parts extracted from bounding boxes around blobs, derived from frame differencing, with CNNs (MobileNet V2^35^). This allows to identify the object independently of its size in the image, since images are rescaled before classification, and was particularly useful for differently-sized animal targets. When image backgrounds were too dynamic for frame differencing (due to motion or exposure changes), we used the FOMO (“Faster Objects, More Objects”) object detection model ^88^ directly on the captured images. The FOMO models may require larger amounts of data for effective training and usually cannot deal well with objects of different relative sizes. All CNNs were trained on the EdgeImpulse platform (https://www.edgeimpulse.com/) to enable their execution on the STM32H7 microcontroller with a closed-source, built-in library.

Portable devices with continuous imaging must balance power consumption and taxonomic resolution. Higher-resolution images invariably require more power, further reducing constrained deployment times. Increasing battery capacity in turn increases cost, bulk - which can restrict mounting possibilities (e.g., cameras in trees) - and theft risk, a major problem with camera traps^83^. Embedded CNNs also need to be simpler than their sophisticated counterparts executed on powerful servers, and the sacrificed precision may enable only coarse classification: This is why we lumped similar-looking non-breeding male and female mandarin ducks into one category. However, embedded devices are particularly relevant for pre-filtering data before transfer to more powerful machines for accurate post-processing (e.g., for a species-level classification).

### Case Studies

#### A) Behavioural Ecology: Monitoring bats visiting Durian tree flowers

We set out to study the behavioral foraging ecology of Durian flower visitors at night, between the end of astronomical dusk (approximately 20:35) and its start at dawn (approximately 4:43), by surveying the only available Durian tree of the Xinglong botanical garden (latitude: 18.734212; longitude: 110.196675 degrees), with flowers at a height of 3 to 5 m. The survey started on 2022-05-20 and ended on 2022-06-13, and was only interrupted by a tropical storm and national holidays.

We used F2 aperture, 12 mm focal length lenses without infrared cut-off filter to allow near-infrared wavelengths to reach the image sensor. We used the maximum resolution of the sensor in grayscale mode (2592 × 1944) and used adaptive frame differencing to detect image changes. Flower visits by bats were a rare event, so we experimented with various setups and exposure parameters on different nights to optimise the image quality and detection performance. We eventually used two near-infrared torchlights (M5, Manta Ray) powered by Osram Oslon Black LEDs with an adjustable light cone angle to illuminate the field of view. From the 25th of May onwards, three 3350 mAh Lithium-ion batteries were connected in parallel inside waterproof battery boxes to power each near-infrared torchlight throughout the night. We tried exposure times between 20 and 50 ms and a gain of 6 to 20 dB to effectively freeze the flying animals’ motion. We detected blobs based on lower pixel area thresholds ranging between 6 000 and 50 000 pixels depending on the object size on the image, maximum pixel area thresholds between 400 000 and 2 000 000, and luminance thresholds luminance values ranging from 3 to 10. The maximum pixel area threshold was often too low: bats would often lower entire tree branches when they hung on the flowers, leading to blobs so large that they were excluded by the maximum area threshold, so we analysed the performance of the blob detection algorithm based on the last five deployments only (9th of June onwards) which had the highest maximum blob area thresholds of 2 000 000 pixels (and a minimum area threshold of 15 000 pixels), a conservative colour threshold of 3, an exposure value of 20 ms, and an image gain of 10 dB.

Based on the last five deployments’ data (during one deployment, the flower did not open), we quantified the accuracy of the detection algorithm by subsetting the detection dataset with variable lower and upper pixel area thresholds to compute the F1 scores at all the value combinations. The total number of false negatives was inferred from their frequency occurrence within saved images (which were image-change triggered), under the assumption that the prevalence of bats was not biased by vegetation movement (rainy or stormy episodes were excluded from deployment periods). Due to the paucity of flowering trees, the short flowering period, and thereby little diversity of images in the training dataset, we were unable to obtain object detection models that were not overfitted for our particular tree. The images resulting from our five last survey deployments were screened visually to log the occurrences of flying or feeding bats, and to assign them to blob detections and the individual flowers they were feeding on. We calculated the average number of visits per flower and plotted the bats’ detections with time to analyse the foraging behavior of the flying fox *Roussettus leschenaultii* in Hainan.

#### B) Landscape ecology: Determining bat and insect occurrence in rice fields

We monitored flying bat and insect activities at night, during astronomical dusk, between approximately 19:20 and 20:30, in a rice field in Hangzhou, Zhejiang province, China (latitude: 30.10247; longitude: 120.05027 degrees). The surveys took place on 6 different nights with an interval of approximately one week, from the 16th of July to the 15th of August 2022. One camera was placed inside the rice field, at a distance of 60 m to its edge, and the other camera was placed at the edge of the rice field, 25 m away from a canal, and 45 m away from the forest edge.

We set up cameras on tripods and pointed them to the sky. We illuminated the sky with four near-infrared torchlights (M5, Manta Ray) mounted on accessory clamps. We used 6 mm F2 focal length lenses without near-infrared cut-off filter, and focused them at a fixed distance of 4 m.

To detect flying objects, we used blob detection with a lower pixel area threshold of 2000 and a higher threshold of 2 000 000 pixels, as well as a luminance threshold value of 5. When blobs were detected, we extracted and saved its bounding rectangle. The image classification model was trained on EdgeImpulse with image extracts obtained from 6 training deployments over 4 nights, resulting in a dataset of 487 images (376 for training, 111 for testing) split over 3 categories: bat, insect, and unknown, when images were not clear enough to distinguish bats from insects (usually because of faraway objects). Our best model had a pixel side length of 96 and achieved an accuracy of 84% for training and testing. Inferencing time was 128 ms, peak RAM usage 346.6K. We then carried out the survey deployments during which the extracted bounding rectangles were classified to bats, insects or unknown flying objects by the camera with the trained model. We determined the field accuracy by manually verifying 1254 detections inside 357 images gathered during the survey deployments.

We used the confidence scores corresponding to the highest F1 score to determine the true events - the number of bat and insect flyovers. These numbers were used as response variables in a binomial mixed effects model with the package lme4^89^ (Bayesian models failed to initialise) using bat flyovers as successes and the sum of all flyovers as the number of trials, with the night as a random effect, and camera location as a fixed effect. We computed the probability that the fixed term’s coefficient differs from zero under the null hypothesis as its P-value.

#### C) Agricultural sciences: Quantifying nocturnal insect rice pests

We monitored nocturnal flying insects by setting up two cameras in rice paddy fields located in Hangzhou, Zhejiang province, China (longitude: 30.10247; latitude: 120.05027 degrees). The cameras were 10 m apart and started at the end of civil dusk, for 2 hours, on the 28th of September, 2022 and on the 28th of June, 2023.

The cameras were fitted with a 4.23 mm focal length lens and powered an internal array of nine white LEDs (1.8 W consumption at 180 lm with a 100% duty cycle; 6000 K color temperature) to illuminate quadratic plastic boards (15 cm × 15 cm, 3 mm thickness). Plastic boards were sprayed with several layers of yellow fluorescent paint (Sparvar, Switzerland) that is commonly used to attract insects during the daytime with yellow pan traps. The plastic boards were held by tablet holders to be parallel with the camera sensor plane, and both camera and tablet holder were held by a metal rail that allowed for precise adjustment of a distance of 9-10 cm between the camera lens and the board. The setup was placed on the concrete walls delimiting rice paddy fields to reduce the occurrence of crawling insects relative to flying insects. We pasted transparent window film on the camera window as a diffuser to achieve more even lighting of the board and insects with less reflections. Even without ultraviolet light source, the boards were effectively attracting nocturnal insects. In preliminary tests, cameras could last 8 hours while powering the LED array. We outfitted the camera with a WLAN module and connected it to a portable modem (brand: 蒙旭, model: MiFi, upload speed 50 Mbps, 10 Ah battery) to send detection data to a web server in real-time.

To detect insects landing on the board, we used blob detection with a lower pixel area threshold of 2000 and a higher threshold of 1 000 000 pixels, and color threshold values of 3 for luminance and the absolute of the a and b channels (Lab color space). When blobs were detected, we extracted and saved its bounding squares to preserve the proportions of the detected targets, as the image classification model requires square inputs. During the 2023 deployment, the full image was additionally saved, lowering the frame rate from approximately 0.40 to 0.10 frames per second - we thus excluded it from the frame rate statistic mentioned previously in the results. We used images from nine test deployments over five nights in August and September, resulting in a dataset of 909 images (735 for training, 174 for testing) split over 10 classes: one for plastic board detections due to false positives, and nine morphospecies that could be classified to order or family level. Our best trained model had a 160 pixels side length and achieved an accuracy of 82% for training, and 97% for the testing dataset. Inferencing time was 248 ms, peak RAM usage 1.4M. We then carried out the actual survey deployments during which the extracted bounding squares were additionally classified with the trained model. We determined the field accuracy by manually verifying 2438 detections inside 1075 images gathered during the survey deployments, obtaining an accuracy of 60% for Chironomidae, 93% for Coleoptera, 8% for Curculionidae, 81% and 55% respectively for the first and second morphospecies of Delphacidae, 66% and 40% for respectively the first and second species of Hymenoptera.

#### D) Pollination ecology: Monitoring bees on rapeseed flowers

Cameras were set up to operate during warm daytimes when bees are most active, between 10:00 and 16:00, inside experimental cages containing *Osmia bicornis* nests and rapeseed plants. We deployed three to four cameras twice, resulting in seven deployments lasting three to four days, in Fuyang district, in an agricultural field (latitude: 30.141028; longitude: 119.937658 degrees).

The cameras were set up on tripods and aimed at the terminal rapeseed flower stands. We used 12 mm lenses with a F2 aperture, resulting in a working distance of approximately 20 cm. We chose flower stands with multiple open flowers and new flower buds, but after three days, the flower stands had sometimes outgrown the picture frame. We set focus as to have the flowers facing the cameras in focus, but the depth-of-field was generally large enough to get acceptable sharpness on the flowers facing away from the camera, behind the flower stand. We only used ambient lighting and adjusted exposure automatically.

The cameras captured pictures at a square resolution (required for object detection models) of 1944 × 1944 pixels, in RGB565, and each image was analysed with an ImageNet v2-based object detection model^88^. Prior to the deployments, we used the same settings for capturing rapes<eed flower images with visiting bees at noon and in the afternoon (estimated total working time: 2 hours) for constituting a training image dataset. We used handheld versions of our camera, consisting of the core H7+ board connected to a powerbank via a USB cable with a flip switch. We selected images that were typical of our deployments and labelled them for bees on the EdgeImpulse platform. The best object detection model had a side resolution of 160 pixels and was trained using 278 images and its performance tested with 81 images, resulting in training and testing accuracies of respectively 95% and 90%. Inference time was 4 ms, peak RAM usage 630.9K. During the deployments, we only saved images with bee detection probabilities above 0.5. We visually screened the first 10 000 pictures of each deployment to determine for each visible (i.e. at least half of the body visible and in focus) bee whether it was detected or not. Based on this, we measured field accuracies ranging from 55 to 89 %, but excluded a stationary false positive (similarly to previous studies^61^) inside one deployment that would otherwise have had a field accuracy of 13%. The stationary false positive was caused by a feature of the exclosure net that triggered the object detection model. The field accuracy may have been low due to the absence of rear-facing bees in the training images, which were collected with handheld cameras that could only be pointed to flowers after bees had approached them.

We extracted hourly temperature data^90^ for Fuyang district to investigate daily activity patterns of bees in relation with the temperature. We constructed a Bayesian negative binomial model using the brms package^91^ and used the number of detections as a response variable, the temperature as a predictor, the cage row as a random effect (as humidity gradients were observed from the left and right sides of the field prior to the study) and a smoothing term based on the time variable, expressed as seconds since midnight, to account for temporal autocorrelation.

#### E) Conservation Biology: Protected waterbird real-time alerts

We installed a camera on the shore of an environmental protection area (Historical and Cultural Reserve) of the West Lake of Hangzhou (latitude: 30.253183; longitude: 120.141125 degrees), Zhejiang province, China, to monitor mandarin ducks swimming in the water, from 7:00 to 13:00. We monitored the mandarin ducks during the breeding season, when males show a striking and distinctive plumage compared to the females.

The camera was placed on a tripod, at an elevated shore position to be able to monitor the water near the shore with a field of view of approximately 1.2 x 1.2 m with a 12 mm focal length, F2 aperture lens. The lake is frequented by passers-by who would feed waterbirds and thus occasionally attract them to the shore. We outfitted the camera with a WLAN module and connected it to a portable modem (brand: 蒙旭, model: MiFi, upload speed 50 Mbps, 10 Ah battery) to send real-time image and detection data to a web server.

The cameras captured pictures at a full resolution of 2592 × 1944 pixels in RGB565. Prior to the deployments, we used the same image settings for capturing images of mandarin ducks of both sexes (estimated working time: 4 half-days) from different shore positions, using the handheld camera setup described above (use case D). We selected images that were typical of our deployments and labelled them for male and female mandarin ducks on the EdgeImpulse platform. The best object detection model had a side resolution of 96 pixels and was trained using 425 images and tested with 87 images, resulting in training and testing accuracies of respectively 93% and 95%. Inference time was 3 ms and peak RAM usage 244.1K. During the deployments, we only saved images with waterbird detection probabilities above 0.1. However, since the original, rectangular images did not fit the required square aspect ratio of the object detection model due to wrong deployment settings, we had to crop the images to square dimensions of 1944 × 1944 pixels in post-processing, using the openMV boards to preserve the original color space that the models were trained with. The loss in image quality resulting from this additional JPEG compression is negligible due to the strong downsampling to 96 side pixels that occurs to feed the images into the model. We then re-processed each cropped image by executing the same ImageNet v2-based object detection model^88^ on them as used during the deployment, to be able to analyse the resulting detections data as follows. We visually screened the first 500 pictures of the deployment to determine for each clearly visible (i.e. at least half of the body visible) duck whether it was correctly detected and identified. 181 images had detections, of which 140 were clearly visible and used to measure field accuracies of 74% for female and 96% for breeding male mandarin ducks.

#### F) Phenology: Flowering plants

We monitored Carthusian pink flowers growing on a planted green space in our university campus (latitude: 30.33406; longitude: 120.03457 degrees) in Hangzhou city, Zhejiang province, China with two cameras, from 2022-09-21 to 2022-11-18. Monitoring was conducted from the start of civil twilight in the morning until the end of nautical twilight in the evening, and interrupted at night, when Carthusian pink closes its flowers.

Cameras were pointed to the ground to monitor an area of approximately 25 × 25 cm with 4.23 mm, F2.8 aperture lenses. The cameras were mounted on tripods with angled metal plates, and a 5 W solar panel, recharging the camera’s internal batteries, was fixed on the tripod in a counter-balancing position to avoid shading the observed flowers. The cameras were set up to verify battery level, auto-expose the scene, take a picture, and execute the object detection model every 15 minutes. In-between the active flower monitoring periods, the cameras went into deep sleep (current consumption: 40 mW) to save battery power, and awoke with an event from the internal real-time clock. In case the battery level dropped below 2.8 V, the camera was programmed to go into a deep sleep mode with regular wakeups at 30 minute intervals to check whether the monitoring could be resumed if the voltage was restored above the threshold by the solar charging, but this situation did not occur. For verifying the images and evaluating the model performance, the images and detection data were transferred to a computer through a USB cable every week during the deep sleep phases.

The cameras captured pictures at a square resolution (required for object detection models) of 1944 × 1944 pixels, in RGB565, and each image was analysed with an ImageNet v2-based object detection model. Prior to the deployments, we used the same settings for capturing unique images of flowers at different spots, at different daytimes and during variable weather conditions (sunrise, sunny noon, rainy noon, evening) to represent the variety of conditions that the object detection model needs to deal with, and we made sure to include increasingly prevalent dead leaves lying on the ground. We labelled images for Carthusian pink flowers on the EdgeImpulse platform. The best object detection model had a side resolution of 320 pixels; it was trained using 118 images and its performance tested with 31 images, resulting in training and testing accuracies of respectively 95% and 90%. Inference time was 12 ms, peak RAM usage 2.4M. During the deployments, we saved all images. We visually screened the first 50 pictures of each camera to determine for each visible flower whether it was detected or not. Based on this, we obtained mean maximum field accuracies of 96%.

To track single flowers, we clustered detections in space and time. We considered only detections above the optimal confidence thresholds of each camera. As object detection sizes and locations could vary even for the same flower, we defined a minimum distance between neighboring object detection centroids of 100 pixels to assign them all to the same day cluster, and then clustered day clusters over days. We defined a minimum lifespan of eight hours to consider day clusters to be flowers, and discarded clusters where less than two thirds of the images had a detection.

## Extended data

**Figure S1:**
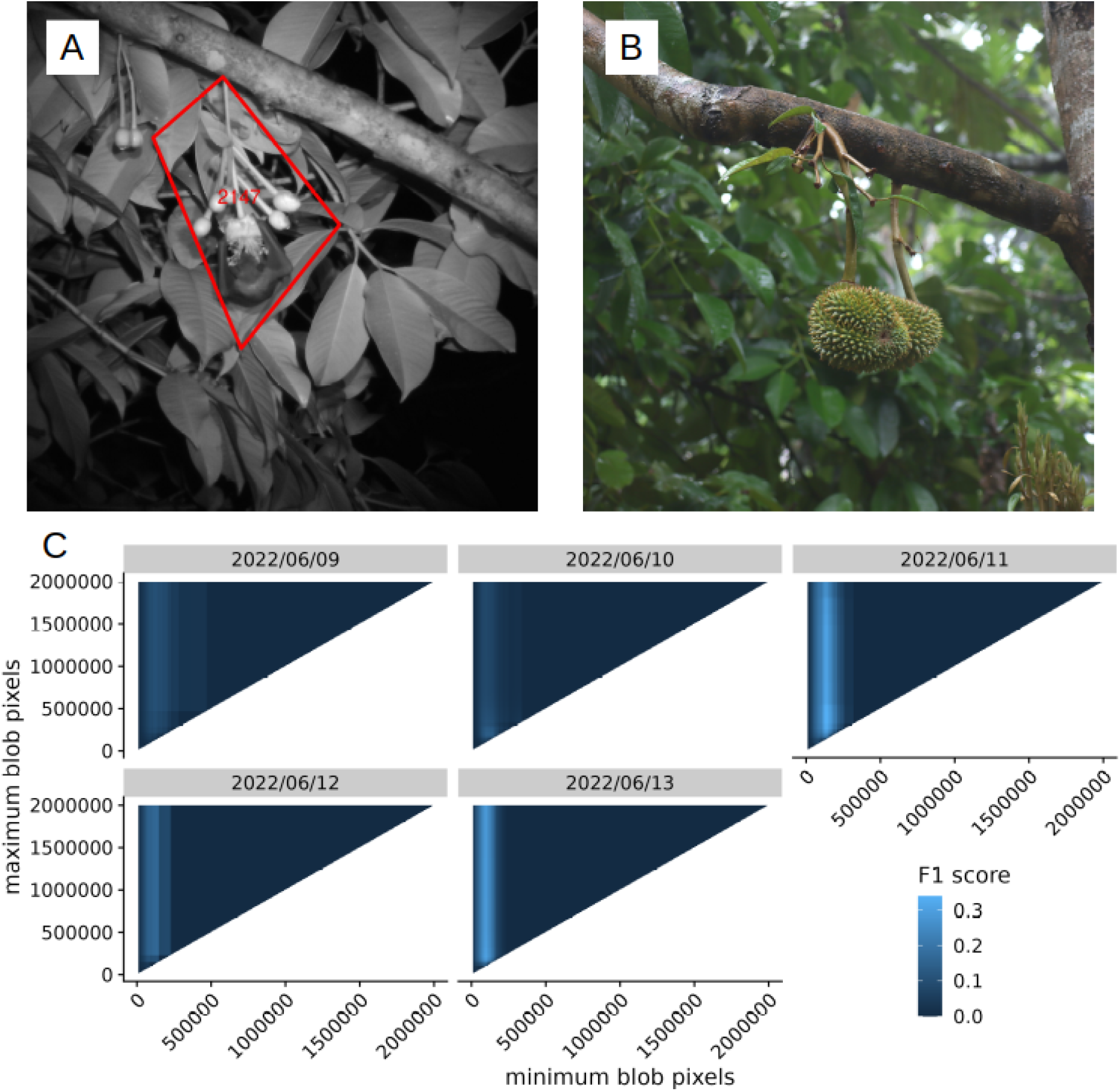
A: a flying fox (*Roussettus leschenaultii*) feeding on a Durian flower, with four outer blob corners drawn as a polygon, and labeled blob ID. B: Durian fruits resulting from pollinated flowers on the same tree. C: Accuracy of the blob detection algorithm for different minimum and maximum blob pixel values on the five different deployment nights.

**Figure S2:**
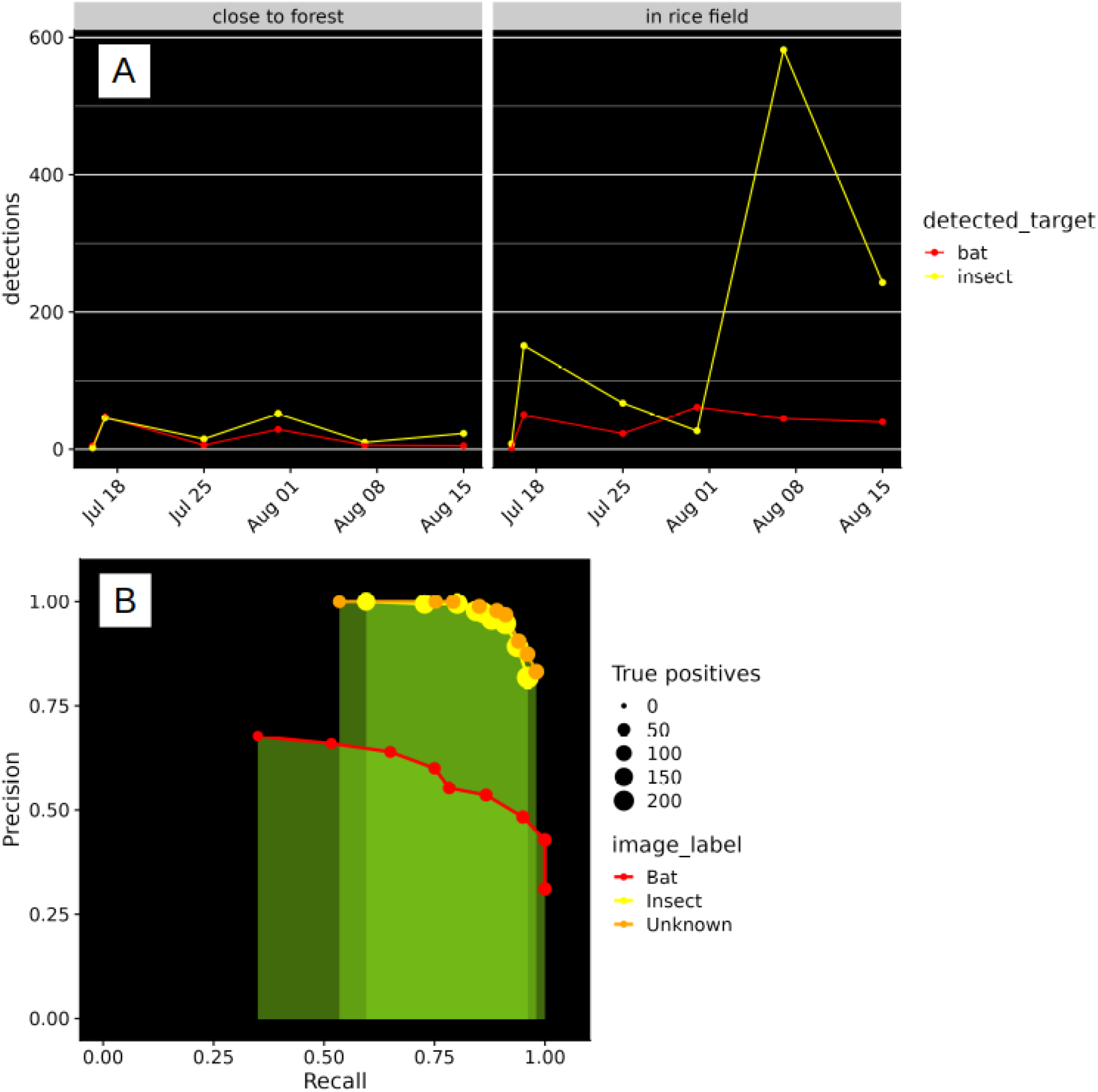
A: Number of detections during each survey night (at maximum accuracy level), drawn for bats and insects and separated by location. B: Precision-recall curves for each detected class based on actual survey deployments.

**Figure S3:**
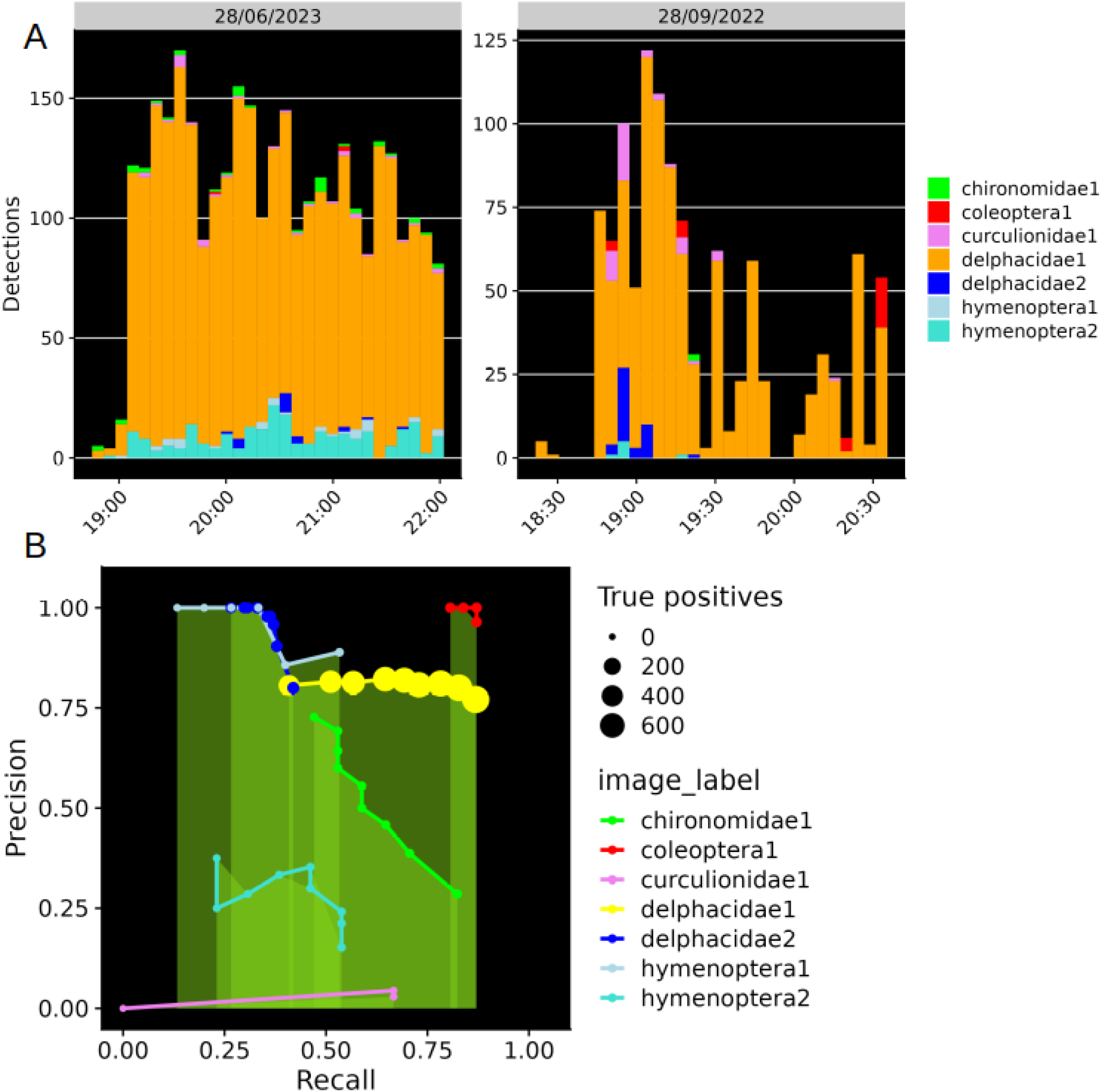
A: Detection frequencies with time per detected class on each deployment night. B: Precision-recall curves for each detected class based on actual survey deployments.

**Figure S4:**
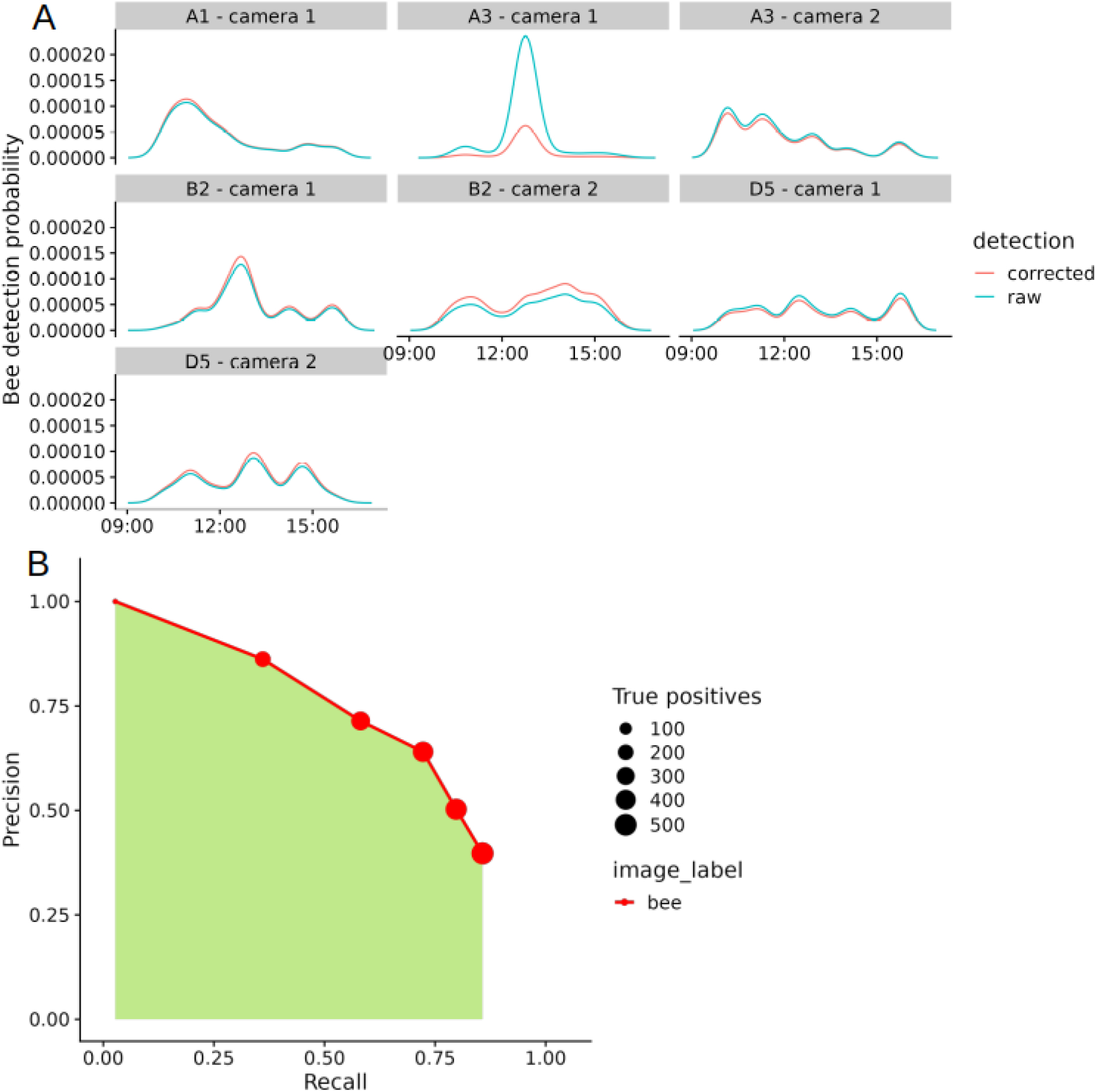
A: Probability density functions for bee detection occurrence with time of the day, separated by deployments. Raw and corrected (standardised) probabilities are shown with different line colors. B: Precision-recall curve for solitary bees (*Osmia bicornis*) based on actual survey deployments, excluding one stationary detection in deployment A3 - camera 1.

**Figure S5:**
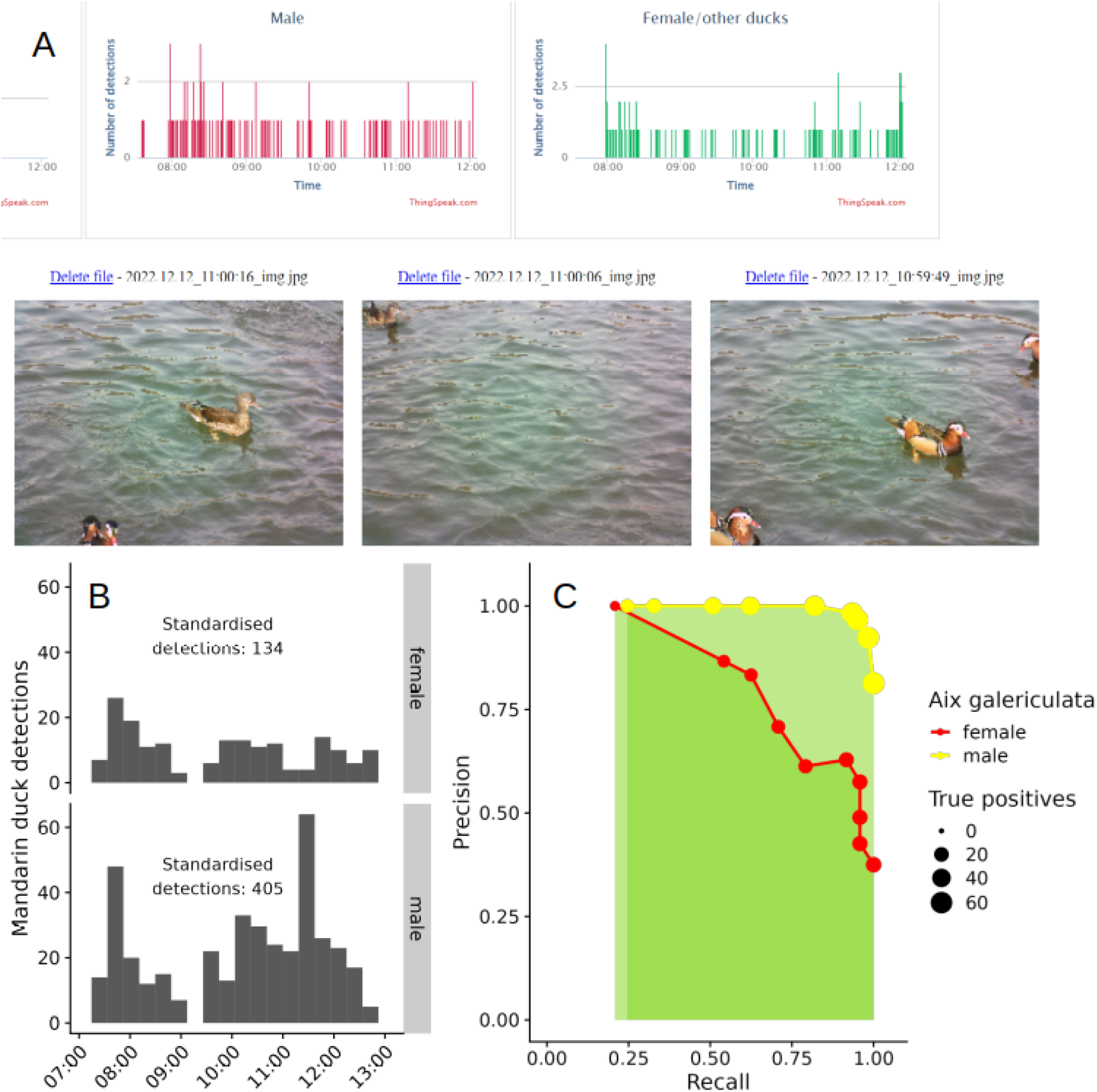
A: Screenshot of a cloud server graphical interface during the deployment, showing transferred images with detected mandarin ducks and bar plots showing the number of detections for each class per time step. B: Detections frequency with time, separated by identified duck sex, and labeled total standardised detections count. C: Precision-recall curve for each detected class based on actual survey deployments.

**Figure S6:**
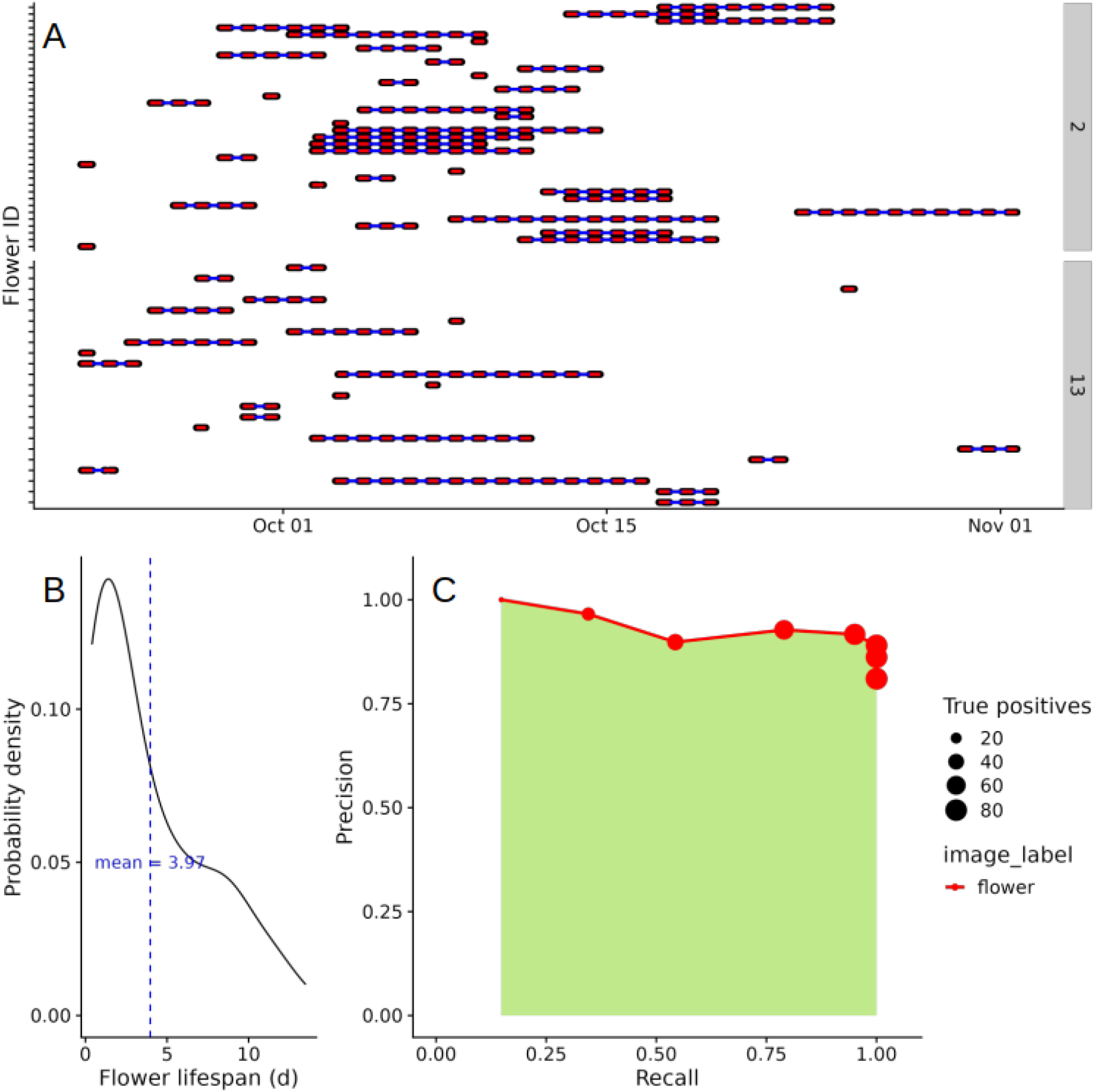
A: Individual tracked flowers (flower IDs omitted from Y axis for clarity), separated by camera (IDs 2 and 13). Red lines represent tracked flowers over single days, and blue lines connect identical flowers through nights. B: Probability density of the flower lifespan; mean shown with a blue dashed vertical line. C: Precision-recall curve for Carthusian pink (*Dianthus carthusianorum*) flowers based on actual survey deployments.

## Data availability

All R scripts and CSV data needed to reproduce the results in the manuscript will be provided on Dryad or a similar repository upon acceptance.

## Acknowledgements

KD thanks his family for all the patience and help with assembling cameras and analysing data. Sincere thanks to Ellena F Yusti, who supported the flying fox identification.

## Author contributions (CRediT)

Kevin Darras: Conceptualization, Data Curation, Formal Analysis, Investigation, Methodology, Project Administration, Software, Supervision, Validation, Visualization, Writing – Original Draft Preparation, Writing – Review & Editing

Marcel Balle: Data Curation, Investigation, Methodology, Software, Validation, Visualization, Writing – Review & Editing

Wenxiu Xu: Data Curation, Investigation, Validation, Writing – Review & Editing

Yang Yan: Formal Analysis, Investigation, Methodology, Software, Validation, Writing – Review & Editing

Vincent Gbouna Zakka: Investigation, Methodology, Software, Writing – Review & Editing

Manuel Toledo-Hernández: Conceptualization, Visualization, Writing – Review & Editing

Dong Sheng: Data Curation, Formal Analysis, Investigation, Validation, Writing – Review & Editing

Wei Lin: Data curation, Investigation, Validation, Writing – Review & Editing

Boyu Zhang: Investigation, Resources

Zhenzhong Lan: Methodology, Supervision

Li Fupeng: Project Administration, Resources, Supervision

Thomas Cherico Wanger: Conceptualization, Funding Acquisition, Investigation, Project Administration, Resources, Supervision, Writing – Original Draft Preparation, Writing – Review & Editing

## Competing interests

The authors declare to have no competing interests.

## Notes

### Competing Interest Statement

The authors have declared no competing interest.

